# ANIMA: Association Network Integration for Multiscale Analysis

**DOI:** 10.1101/257642

**Authors:** Armin Deffur, Robert J Wilkinson, Bongani M Mayosi, Nicola Mulder

**Author notes:** Corresponding author at: Department of Medicine, Old Main Building, Faculty of Health Sciences, University of Cape Town, Anzio Road, Observatory, 7925, South Africa.

## Abstract

Contextual functional interpretation of -omics data derived from clinical samples is a classical and difficult problem in computational systems biology. The measurement of thousands of datapoints on single samples has become routine but relating ‘big data’ datasets to the complexities of human pathobiology is an area of ongoing research. Complicating this is the fact that many publically available datasets use bulk transcriptomics data from complex tissues like blood. The most prevalent analytic approaches derive molecular ‘signatures’ of disease states or apply modular analysis frameworks to the data. Here we show, using a network-based data integration method using clinical phenotype and microarray data as inputs, that we can reconstruct multiple features (or endophenotypes) of disease states at various scales of organization, from transcript abundance patterns of individual genes through co-expression patterns of groups of genes to patterns of cellular behavior in whole blood samples, both in single experiments as well as in a meta-analysis of multiple datasets.

## Introduction

The human immune system can be regarded as a complex adaptive system^1^, even though it is integrated with the more complex system of the whole organism. A true systems view^2^ of the immune system needs to account for the various aspects that characterize complex adaptive systems generally. This includes emergence, non-linearity, self-organization, noise, scaling, heterogeneity, a network architecture and preservation of context for individual observations^3^. Whole blood is a “window” onto the immune system^4^, allowing a reasonably detailed assessment of the overall state of the immune system based on analysis of easily obtained blood samples. Human diseases can be conceived of as complex state vectors embedded in high dimensional state space.

Here we present ANIMA, Association Network Integration for Multiscale Analysis, a framework for discovery of high order, high complexity and low dimensional state space vectors from high dimensional low order, low complexity data (i.e. non-normalised microarray data and clinical/ sample phenotype data) by producing and interrogating a multiscale association network, which allows summary and visualization of different, but simultaneously valid views of the state of the immune system under different conditions and at multiple scales.

To showcase this, we have chosen three publically available datasets which were generated using whole blood from human subjects with one of three infectious diseases (acute HIV infection, malaria and respiratory viral infections).

## Method outline

*ANIMA* generates multiscale association network components from multiple data types (expression data, clinical data and annotation data, e.g. biological pathways databases), and merges all components into a single network (Figure 1).

**Figure 1.**
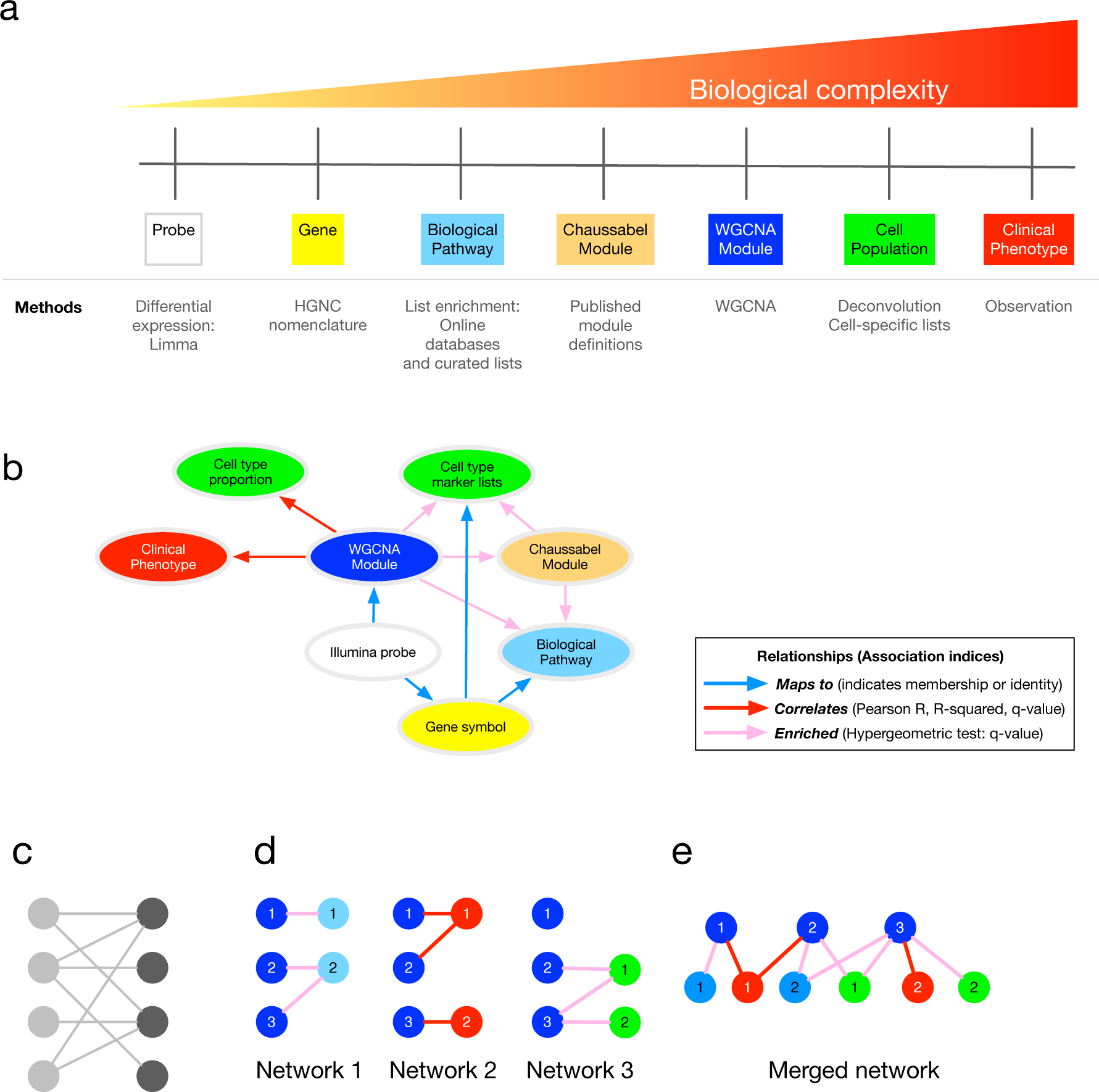
Method overview. (A) Analytical approaches and biological complexity. (B) Relationships between output types (C) A bipartite graph, with two classes of nodes connected by edges. (D) The separate bipartite graphs, with one node type in common. (E) Multipartite graph obtained after merging the three graphs in (D). Abbreviations: HGNC, HUGO Gene Nomenclature Committee; WGCNA, weighted gene co-expression network analysis.

First, the algorithm constructs twenty-nine bipartite networks from a multi-scale analytic pipeline (Figure 1A, C), combining the output, and the relationships between different classes of output, of various analytic approaches (Table 1). Each network contains the associations between two distinct data types (Figure 1B, D, Table 2). The final association network is a result of graph union, merging all bipartite networks on shared node types while retaining all edges (Figure 1E). This results in a data structure that exposes the relationships between modular gene expression and higher-level phenomena, while retaining key probe- and gene-level information. Three types of association (with their respective association indices) are utilised, resulting in three distinct edge types in the final data structure (See Figure 1B, 1E, Table 2 and Online Methods). Weighted gene co-expression network analysis^5^ (WGCNA) is the core analytic method in ANIMA as this is used to discover biological processes in the system of interest.

**Table 1.** List of analytic approaches used in constructing the ANIMA database.

| Approach | Method | R package/<br>implementation | Reference |
| --- | --- | --- | --- |
| Array data import and normalisation | variance stabilising transformation, quantile normalisation | lumi | 16 |
| Probe filter | quality filter to remove non-informative probes from analysis | ReMoat, ReAnnotator | 17,18 |
| Differential expression | linear model/ moderated t-test | limma | 19 |
| Estimate of cell-type proportions | Deconvolution by least-squares fitting | CellMix | 20 |
| Chaussabel module expression | Published module definitions and custom R code |  | 11,14 |
| Chaussabel module differential expression | linear model/ moderated t-test | limma | 19 |
| WGCNA module detection | Clustering of topological overlap matrix and dynamic tree cutting | WGCNA | 5 |
| Chaussabel and WGCNA module annotation | List enrichment testing by hypergeometric test and multiple testing correction | WGCNA (UserListEnrichment); ReactomePA | 5,21 |
| WGCNA module metrics | Differential expression, module AUC, signature enrichment | Custom code |  |
| Module eigengene correlations | Pearson or Spearman rank correlation | WGCNA, custom code | 5 |
| Construction of bipartite graphs | Bipartite graphs from adjacency lists (based on list enrichment) or incidence matrices (based on correlation) | igraph | 9 |
| Merging of all bipartite graphs | Graph union | Neo4j: cypher command *merge* | 22 |

**Table 2.** Types of associations used in the ANIMA database

| Association type | Association index | Intermediate result | Multiple testing correction |
| --- | --- | --- | --- |
| Correlation | Pearson R or Spearman rho | Incidence matrix | Yes |
| List enrichment | hypergeometric index | Adjacency list | Yes |
| Mapping | simple mapping | Adjacency list | Not applicable |

Network construction combines three processes. Firstly, mathematical operations on data are performed, independent of prior knowledge. This aspect of the approach is completely unsupervised. The second process involves the testing of hypotheses, to determine differential transcript abundance and differential co-expression of transcripts, requiring knowledge of phenotype/ trait classes for the samples. Finally, the results of the first two processes are integrated in various ways with prior knowledge. The result is a collection of statistically robust analytic results and various associations between them and known biology, in the form of a large, multipartite graph. This large graph is stored in a Neo4j graph database (referred to as the ANIMA database), making the overall network structure as well as the individual nodes and edges accessible for further analysis (see Online Methods).

After network construction, information in the graph is accessible and utilized to expose new information not present in any of the individual steps. The key to making the ANIMA database useful lies in the use of functions and web applications (see Online Methods) that query this large multipartite graph and return visualization of relationships or tables of nodes and/ or links (associations between nodes). This has been implemented in several R functions which underpin a Shiny web application called *ANIMA REGO*.

The novelty and value in ANIMA lies in its three broad approaches to data interpretation: Firstly, it allows detailed, multiscale investigation of a single dataset with a focused research question where two phenotype classes are compared (**multiscale class comparison**).

Secondly, as datasets are stratified by default using two variables (typically one identifying two disease classes, and the other either a potential confounder (e.g. sex), or a second biologic variable of interest, (like co-infection with a second pathogen), we can examine the interaction of the second variable with the first using a factorial study design (**factorial analysis**). Finally, multiple datasets, and multiple conditions can be meaningfully compared and contrasted to identify similarities and differences (**meta-analysis**), both at cellular and modular level.

## Results

### Validation study: data sets

We analysed three publically available microarray datasets using the ANIMA pipeline and toolset (ArrayExpress/ GEO identifiers: E-GEOD-29429, E-GEOD-34404/ GSE34404, E-GEOD-68310/ GSE68310). The first compares a cohort of subjects with acute HIV infection to healthy controls^6^, the second compares symptomatic malaria to asymptomatic controls in children from a malaria-endemic region^7^, and the last compares the host response in early symptomatic viral respiratory infections^8^. Where the datasets contained samples from multiple timepoints, we restricted the analysis to healthy controls and the first disease timepoint. The respiratory virus infection dataset contained samples from subjects with infections other than influenza or rhinovirus; we excluded those from this analysis. **Figure S1** shows the experimental design for the factorial analysis in *limma* for each of the datasets, together with the numbers of samples in each of the individual groups. Crucially, each of the datasets also included clinical data, which was integrated in the analysis.

### The ANIMA network

The result of the script that builds the ANIMA network is a large, mostly connected, multipartite graph. **Figure S2** shows a subgraph of this network (for dataset **HIVsetB**, edge 5).

### Searching the ANIMA network using network paths and filtering on node and edge properties

In the most direct approach, the large ANIMA graph in the neo4j database can be searched directly from a web browser using the Cypher Query Language (CQL) (https://neo4j.com/developer/cypher-query-language/).

For instance, the query

~~~
MATCH (ph:pheno)-[r1]-(n:wgcna {square:'HIVsetB',edge:5})-[r2]-(p:PROBE)-[r3]-(s:SYMBOL) WHERE r1.weight > 0.6 AND p.logfc > 2 RETURN *
~~~

returns WGCNA modules for the HIV positive vs control comparison in the **HIVsetB** dataset whose module eigengene (ME) is strongly correlated with a clinical variable (Pearson R > .6), and it also returns the genes mapping to probes within those modules that have high levels of over-abundance in HIV-1 infected individuals. In addition to the standard web browser interface for neo4j, into which the above query can be directly entered and returned in the browser window (Figure 2A), we also provide a function in R (`igraph_plotter`) that returns the network found, and plots this within R (Figure 2B), exports the network as node and edge lists for import in other software like Cytoscape (Figure 2C), or returns the result as an *igraph*^9^ object for further manipulation within R.

**Figure 2.**
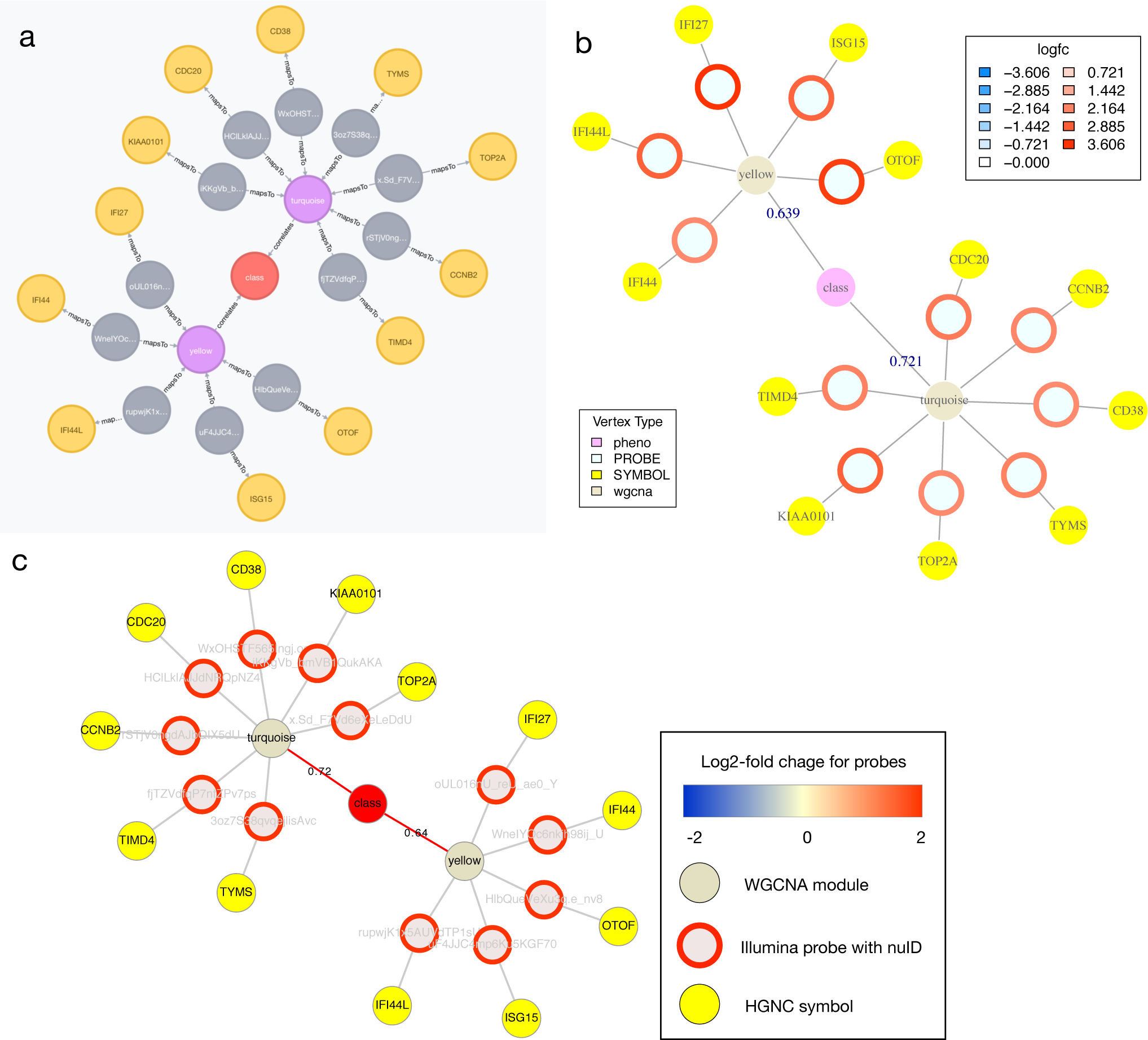
Visualising Cypher query results. Relationships between nodes extracted from the ANIMA database using a Cypher query applied to the **HIVsetB** data (N_HIV_=30, N_Controls_=17, see **Figure S1**). Shown are two WGCNA modules that contain probes with increased transcript abundance in acute HIV infection and whose module eigengene is positively correlated with disease class (an ordinal variable). (A) Result from native browser interface for Neo4j. (B) Result plotted from within an R session connected to the ANIMA database, using the *igraph_plotter* function. Log2-fold change values for the individual probes are shown by coloured rings; values are shown in the legend. (C) The same result, visualized in Cytoscape, taking advantage of the *igraph_plotter* function to export node and edge lists for easy import into Cytoscape. Links/ edges are annotated with Pearson correlation coefficients where applicable.

### Approach 1: Multiscale class comparison

#### Accessing individual transcript abundance levels in multiple conditions

It is useful to view transcript abundance patterns of specified probesets, for instance to compare microarray data to RT-PCR validation data, or to investigate the behavior of groups of biologically related genes in various conditions. We provide an interface for this in ANIMA. The user can submit a search string (in the form of a regular expression) containing gene names (HUGO Gene Nomenclature Committee (HGNC) symbols), and box-and whisker plots for the results are returned. In the original paper on acute HIV infection the authors discuss a gene set of six conserved genes that appear at multiple timepoints in an inferred regulatory network of viral set point^6^. We show the normalized expression data stratified by HIV status and sex in the two datasets included in the HIV analysis for these six genes (Figure 3). In the paper on acute viral respiratory infection, IFI27 and PI3 are identified to differ between acute influenza A and human rhinovirus infections. In influenza, IFI27 is upregulated and PI3 downregulated relative to human rhinovirus. The malaria study replicated prior knowledge of differential transcript abundance for C1QB, MMP9, C3AR1, IL18R and HMOX1; we show similar results for these transcripts. Supplementary Table 1 lists the results of differential expression analysis for the above transcripts in the three conditions, providing validation of data at individual transcript level.

**Figure 3.**
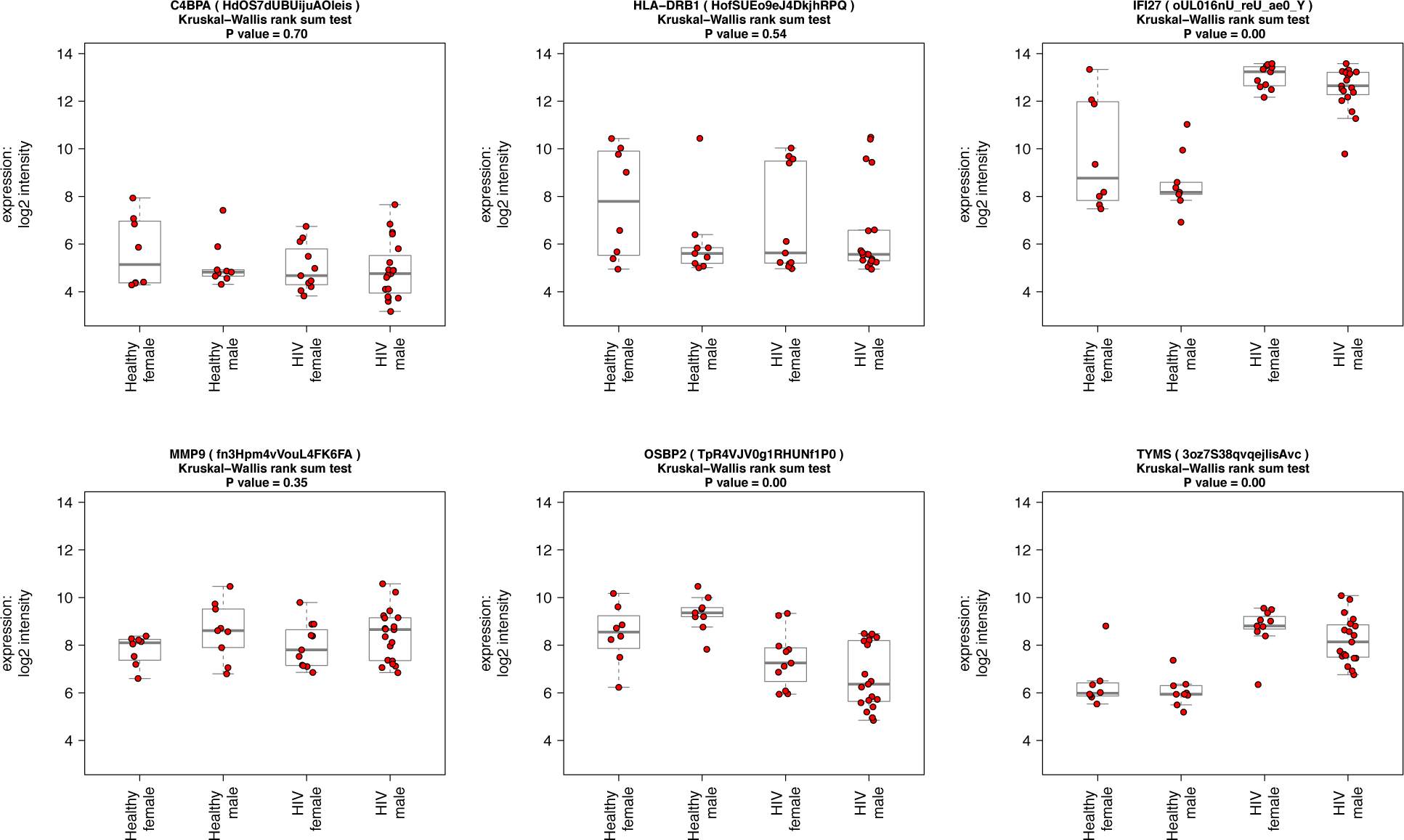
Visualising individual probe-level expression data. Box-and-whisker plots showing normalized, log2-transformed probe-level expression data for six selected genes, obtained by a custom function in R in four groups: Healthy female, N = 8, Healthy male, N = 9, acute HIV female, N = 11, acute HIV male, N = 19; data from **HIVsetB** dataset. Gene (and probe nuIDs for disambiguation) are given for reference; the y-axis shows log2 scale normalized intensity values. Box and whisker plots show median, interquartile range, and range. Outliers are defined as values that lie beyond the whiskers, which extend to maximally 1.5 X the length of the box. Individual datapoints are superimposed in red on the box-and-whisker plots. The four groups are compared using Kruskal Wallis rank sum test and the P-value for the comparison is shown in the plot title. Results for individual pairwise comparisons are not shown.

#### Functional annotation of WGCNA modules

An important question is what do WGCNA modules represent, given that these are groups of genes that co-vary across samples, but that can differ dramatically in size. Instead of searching only for evidence of co-regulation of expression by transcription factors, one should consider other causes of this co-variance. We propose the notion of WGCNA modules representing biological processes that may be regulated at different scales. For instance, a group of transcripts will co-vary across samples if these transcripts are expressed (predominantly) in a single cell type, and the proportions of this cell type varies between samples (Figure 4). Another group of transcripts may be expressed in multiple cell types, but represents a concerted transcriptional program executed in response to a specific stimulus, such as interferon-alpha stimulating the expression of a specific group of interferon-regulated genes (see below). Generally, we determine the function of WGCNA modules based on all statistically significant associations with pathways and cells.

**Figure 4.**
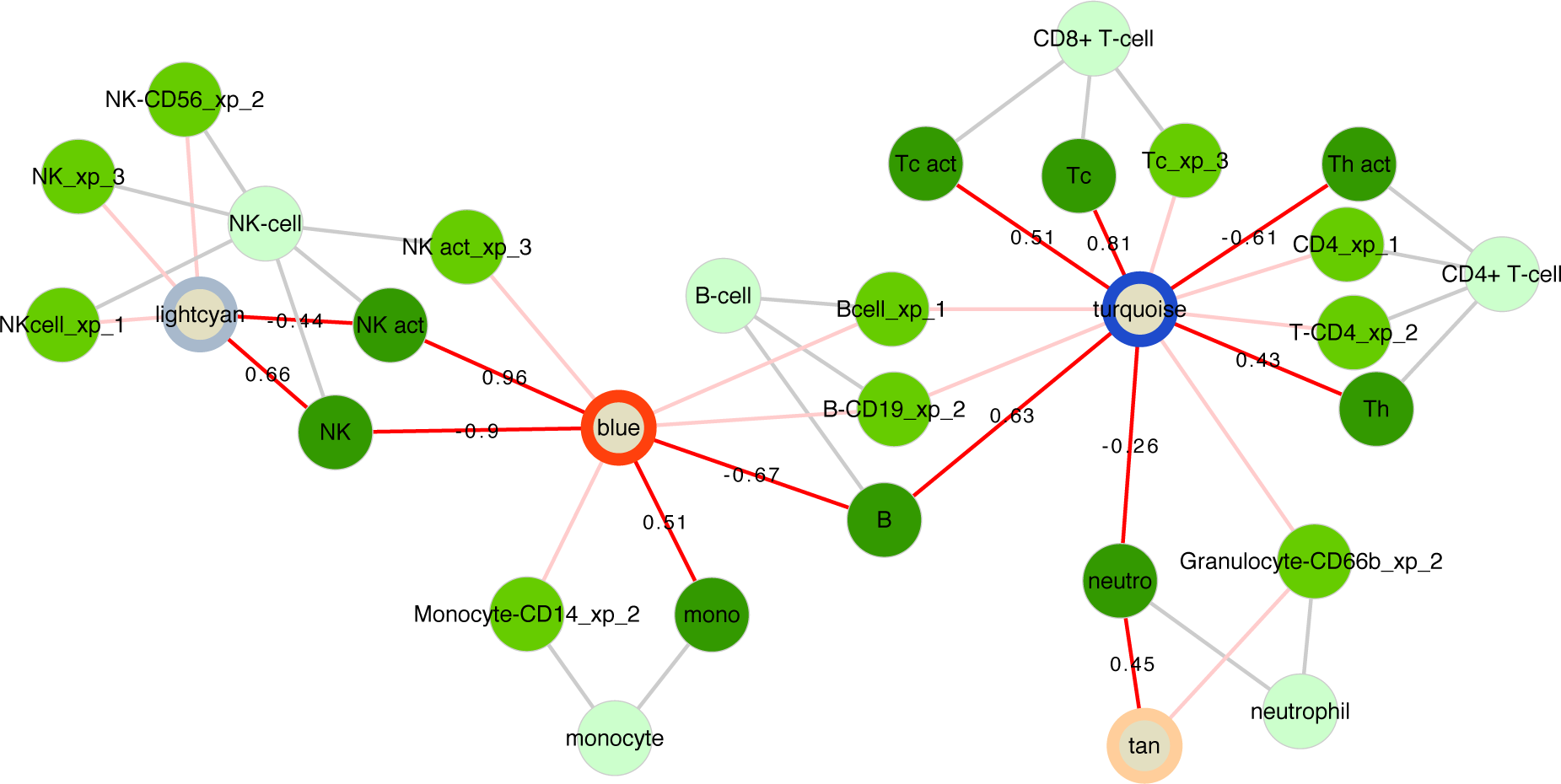
Cell associations of WGCNA modules. Relationships of WGCNA modules and different cell types in the **respInf** dataset (Day 0 acute influenza, N = 46 vs baseline healthy samples, N = 48, see **Figure S1**). Shown are WGCNA modules whose expression correlates with specific cell-type proportions (dark green, edges annotated with Pearson correlation coefficient *R*) *and* that are enriched for the genes specific to that cell type (medium green, suffixes xp_1–3 indicate the respective gene list on which the cell assignments were based, see Online Methods). The classes of cells are indicated in light green. The modules are annotated with coloured rings representing the difference in median eigengene values between cases and controls (diffME, see Online Methods); blue indicates modules which are under-expressed, and red indicates modules that are over-expressed in cases relative to controls.

#### Relationship of modules to clinical variables

We performed Pearson correlation of WGCNA module eigengenes (ME) and clinical variables (described in Online Methods). Figure 5 shows correlation of the *pink* ME with age and CD4 count in the HIVsetA data set. Clearly, the *pink* module is significantly associated with CD4 count, an association that is independent of age. Further investigation shows that this module is linked to interferon signaling, and probably expressed in a variety of cells. This agrees with our understanding of acute HIV infection, which is associated with a robust type-I interferon response and an acute drop in CD4 count^10^.

**Figure 5.**
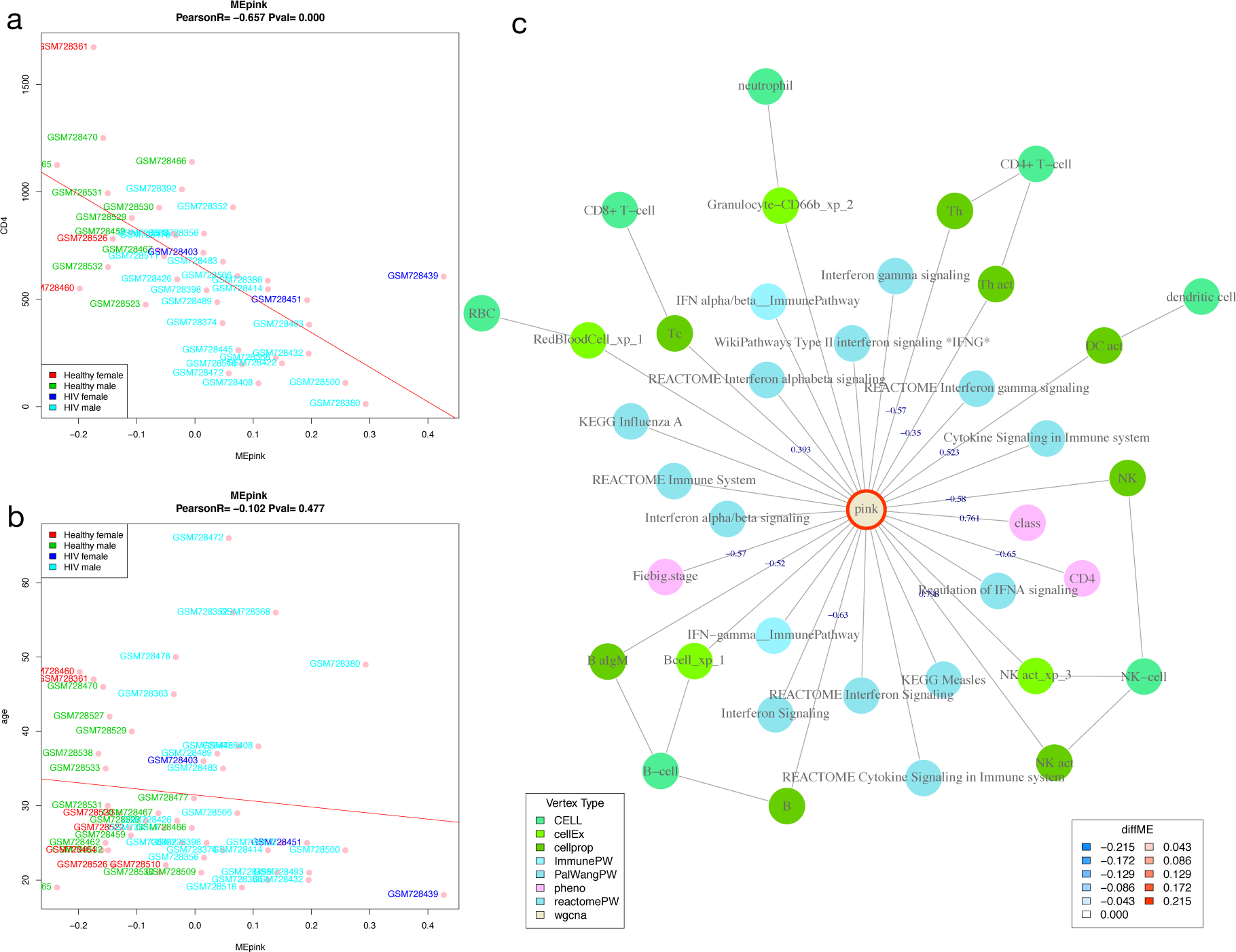
Correlation of module eigengenes with clinical variables. Shown is the Pearson correlation of the *pink* module eigengene with CD4 count (cells/microlitre) (A) and with age (years) (B) in acute HIV (N = 28) vs healthy controls (N = 23) in the **HIVsetA** dataset. Study subject IDs are used as point labels, and coloured as indicated in the legend. Plot titles show the Pearson coefficient R and the associated P-value. (C) WGCNA module annotation obtained from the Neo4j database for the *pink* module. Edges are labelled with the correlation coefficient (R) where applicable. Note that the same coefficient is obtained for CD4 count as in panel A. Legends are shown for vertex type and **diffME** (a measure of differential coexpression (see Online Methods), i.e. the extent that the module eigengene median varies between two classes). Abbreviations: diffME, differential module eigengene.

#### Investigating the structure of WGCNA modules

WGCNA modules are groups of co-expressed transcripts. Demonstrating the extent and direction of correlation of their constituent probes, and their relationships to biological pathways requires sophisticated visualization (Figure 6). We showcase the example using the **HIVsetB** dataset and focus on two modules. The *turquoise* module shows the coordinated action of genes involved in cell division, relating to the cell proliferation in the lymphoid compartment. The *yellow* module is clearly related to interferon signaling; of interest here is that within this module there is a group of transcripts highly correlated with each other, all related to interferon signaling, suggesting that these transcripts may all be downstream of a single regulatory factor. Also clear from the figure is that this module is not limited to a single cell type, but rather to innate immune cells in general.

**Figure 6.**
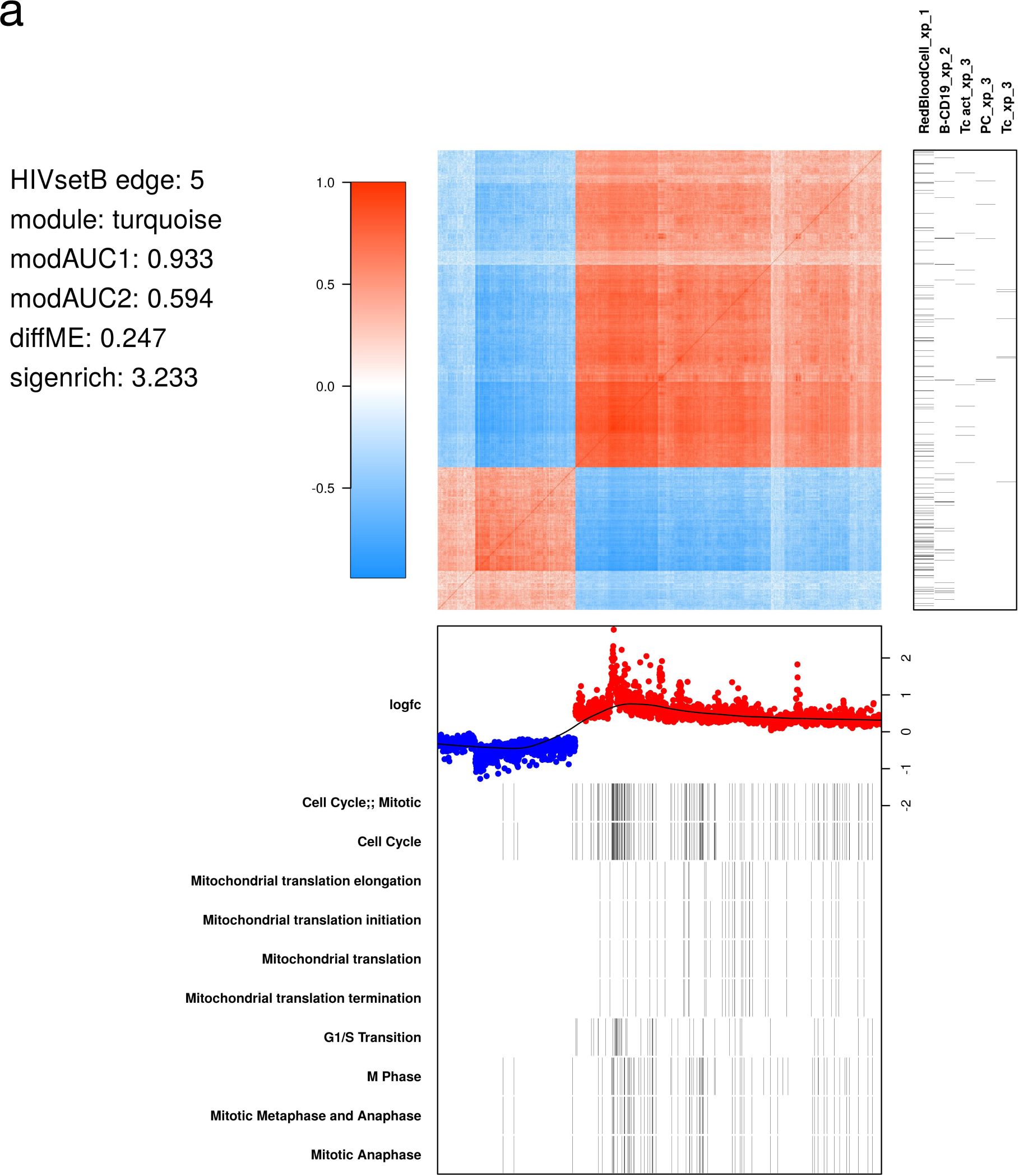
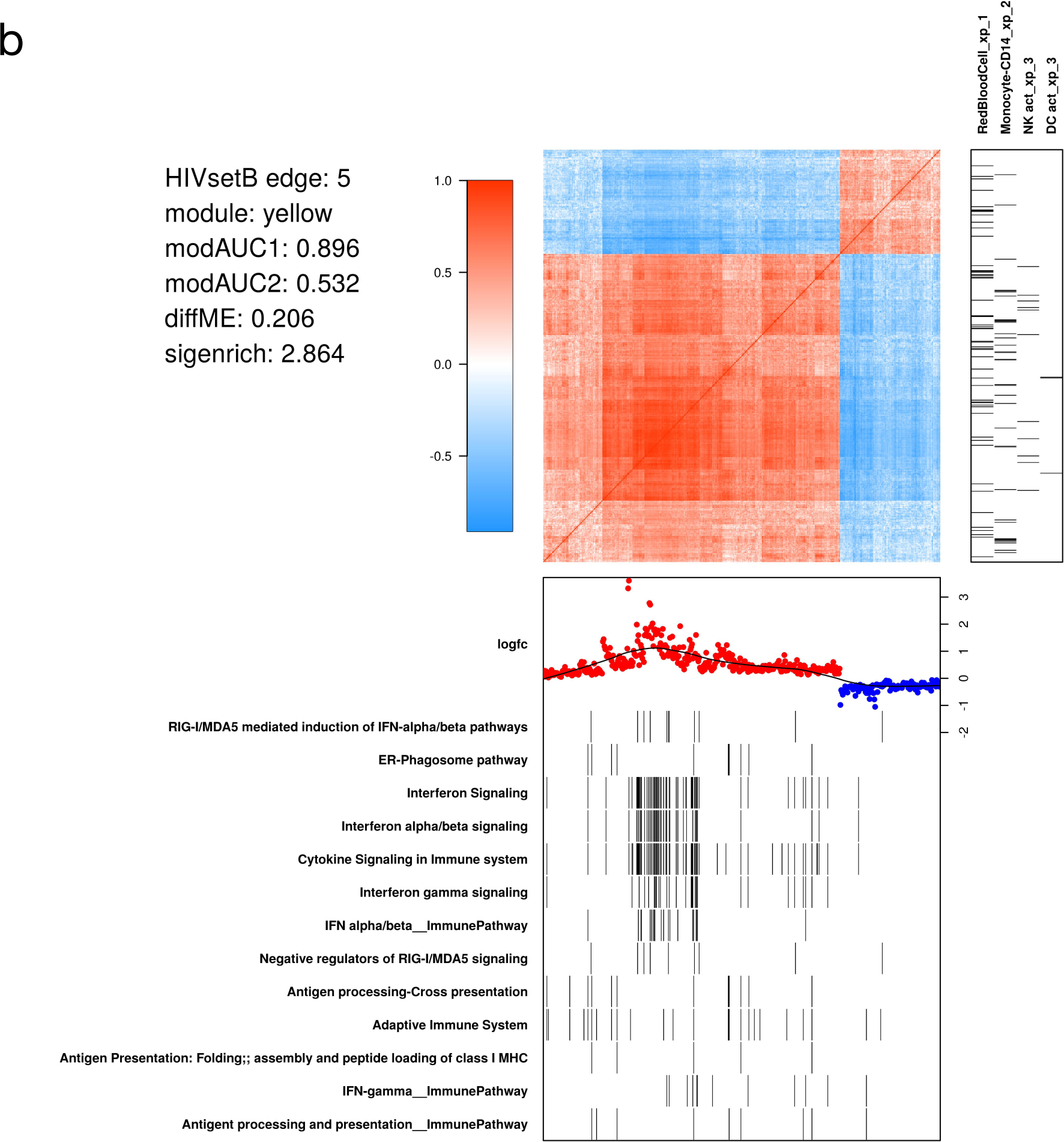
WGCNA module structure. (A) Correlation matrix of all probes in the *turquoise* module in the HIVsetB dataset (N_HIV_=30, N_Controls_=17, see **Figure S1**). Colours in the heatmap represent Pearson correlation coefficients, ranging from −1 to 1, as indicated by the legend. The module is enriched for lymphocyte-specific genes (right annotation panel) as well as cell cycle/ mitosis associated genes, suggesting that various lymphocyte subsets in acute HIV infection are actively proliferating. (bottom annotation panel). Log_2_-fold change values refer to differential transcript abundance in acute HIV relative to healthy controls. (B) Correlation matrix of all probes in the *yellow* module in the HIVsetB dataset. It is enriched for innate cell genes as well as interferon signaling, suggesting that innate immune cells are in an interferon-induced state. Additional annotation information is provided to the left of the heatmap. The parameters *modAUC1, modAUC2, diffME* and *sigenrich* are defined in Online Methods. The plot is generated using a custom R function (*mwat*).

#### Relationships between module-based approaches

Both WGCNA and the modular approach pioneered by Chaussabel, et al^11^ rely on clustering of similarity matrices to derive modules. A key difference in these methods in ANIMA is that the Chaussabel modules are pre-defined, whereas the WGCNA modules are derived from the transcriptional data under study. It is therefore interesting to discover the relationships between these two approaches. Figure 7A shows the bipartite network of WGCNA and Chaussabel modules derived from the **HIVsetA** dataset, as defined by the hypergeometric index. Figure. 7B and 7C show the two projections of the bipartite graph. The hypergeometric index is not the only way associations between the two module types can be demonstrated; for instance, a more indirect association can be inferred when a WGCNA and Chaussabel module map to the same biological pathway. It is clear from the plots that only a subset of the list of Chaussabel modules associates with WGCNA modules in any given condition.

**Figure 7.**
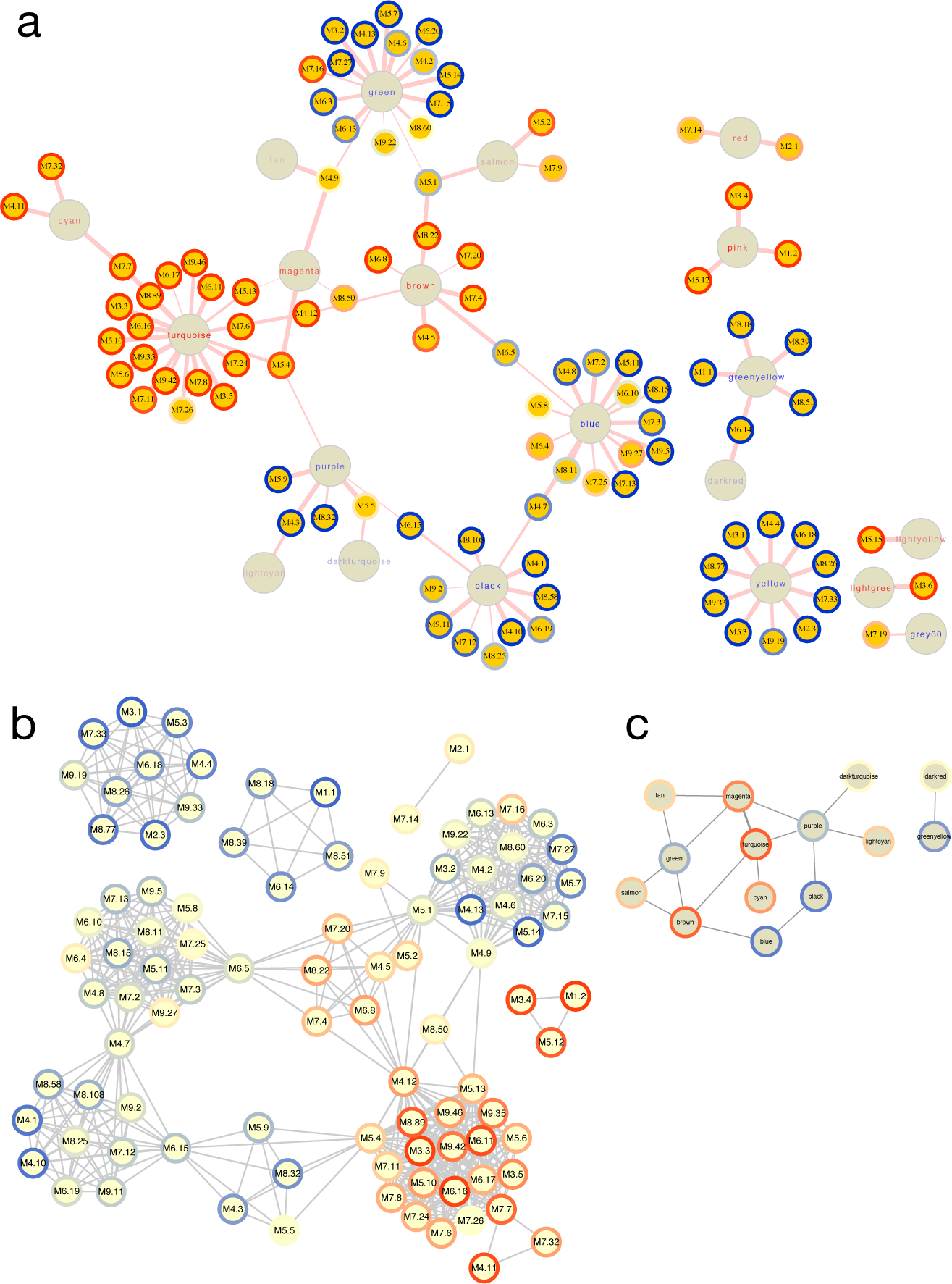
Relationships between WGCNA and Chaussabel modules. (A) Bipartite graph of the two module types based on the hypergeometric association index in the **HIVsetA** dataset (acute HIV, N = 28 vs healthy controls, N = 23). Strikingly, Chaussabel modules tend to have the same direction of differential expression (indicated by the rim colour of the Chaussabel modules, red indicating up-regulation in acute HIV, and blue indicating downregulation) as WGCNA modules they map to, indicated by the label colour of the module. (B) Projection 1 of (A), showing relationships between Chaussabel modules based on shared WGCNA modules; dense cliques of modules are observed. (C) Projection 2 of (A), showing relationships between WGCNA modules based on shared Chaussabel modules. All associations (hypergeometric test) shown are corrected for multiple testing, BH-corrected P-value < 0.05. All outputs were generated using the *igraph_plotter* function, exporting vertex and edge tables of the bipartite graph and the two projections and importing these into Cytoscape.

#### Deconvolution

An important question in analyzing transcription data from complex tissues is whether differences in transcript abundance are attributable to transcriptional regulation in one or more cell types, or to changes in the composition of the overall leukocyte populations.

**Figure S3** shows the results for the **HIVsetB** dataset, and **Table S2** shows the results of non-parametric statistical testing for differences in median cell-type proportions, per cell-type and class comparison with uncorrected P values as well as P-values corrected for multiple testing using the Benjamini-Hochberg procedure; these were encoded as parameters *diffP* and *diffQ* of the *cellprop* node type in the ANIMA database. Neutrophils were the most abundant cell-type estimated from the array data. The proportions of activated NK cells and CD8+ T cells were significantly elevated in acute HIV infection, and proportions of B cells and CD4+ T cells were reduced, in line with published observations^12^.

#### Virtual cells: an estimate of functional phenotypes of different immune cell types

Given that the immune response is mediated by different types of cell, we attempted to recreate “virtual cells” based on the assumption the genes that co-vary in terms of transcript abundance across samples with that of cell-type specific genes are expressed in that particular cell type. With these relationships, we generated virtual cells (probe co-expression matrices annotated with biological pathways and probe differential expression data). **Figure S4** shows two cells (B-cells and neutrophils) in acute HIV infection; both characterized by interferon signaling. Given the “virtual cells” and their functions we can compare pathway-level transcript abundance in different cell types by creating a matrix of cell types and pathways, with each entry representing the “pathway activity” for a given cell and pathway combination. To illustrate this, we show replication of the finding of NK-cell activation in acute influenza infection (**respInf** dataset, Figure 8 A,B), and in addition provide more detail on which pathways are probably up-or downregulated in these and multiple other cells.

**Figure 8.**
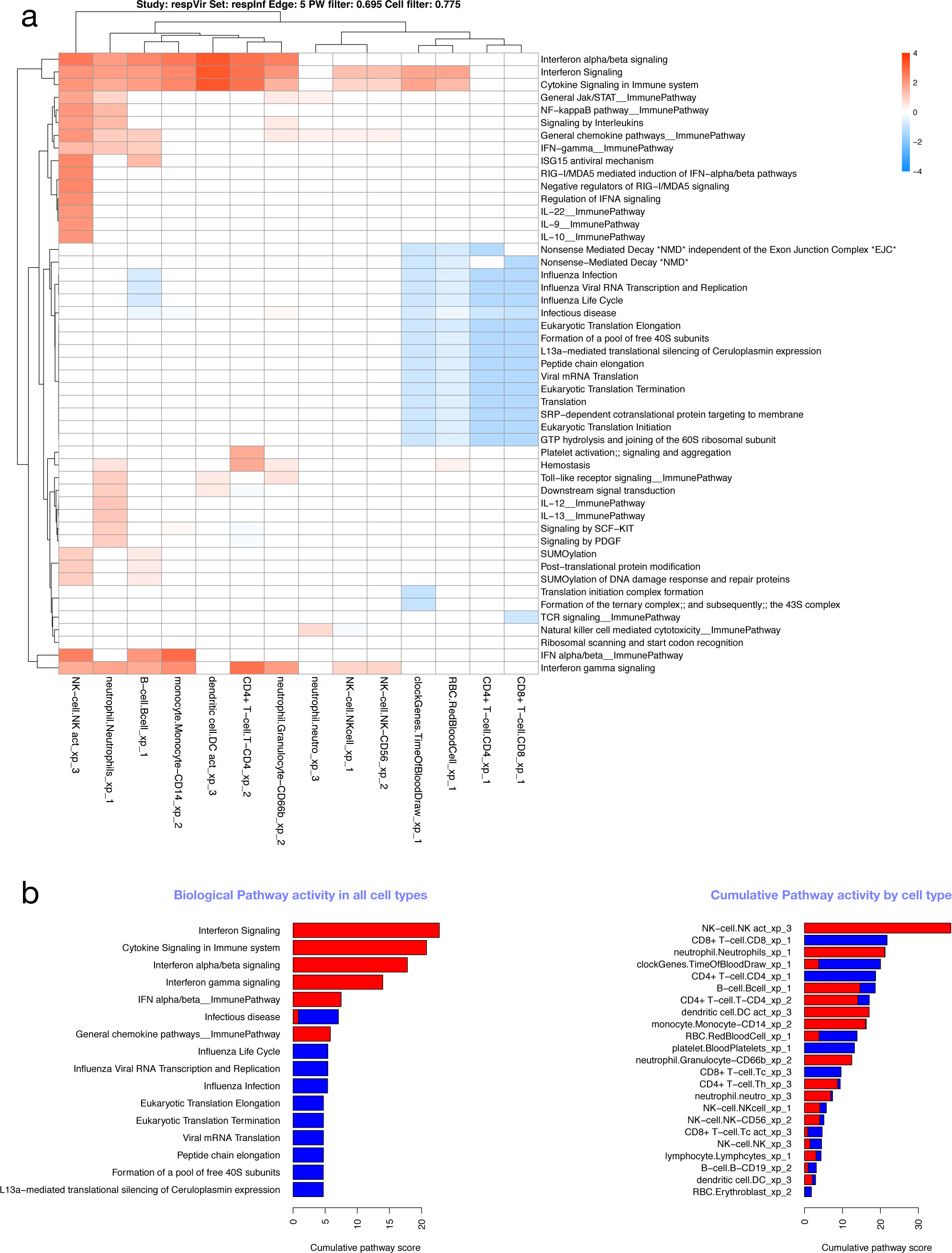
Cell/pathway activity matrix. (A) Cell/pathway activity matrix for all cell-types for the **respInf** dataset (Day 0 acute influenza, N = 46 vs baseline healthy samples, N = 48, see **Figure S1**). The clustered heatmap shows pathway activity scores representing the mean log-2 fold change for all probes in the pathway for a particular cell type (see Online Methods). There is a clear interferon response in multiple cell types, as well as down-regulation of other pathways associated with translation. (B) Barplots highlighting the most highly differentially regulated pathways (left panel, determined by row sums of matrix in A), and cells with highest levels of differential expression (right panel, determined by column sums of matrix in A). In all cases, up- and downregulated pathway scores are kept separate.

### Approach 2: Factorial analysis

#### Modules driven by sample class or sex

Our main interest in investigating transcriptomic datasets is to identify molecular and cellular processes that drive, or at least are associated with, specific phenotypic traits of the samples. Therefore, the experimental design for the differential abundance analysis and the probe filtering steps prior to WGCNA module discovery were both designed to highlight processes associated with two factors: disease class and sex; the former because this is the basis of the research question for the three studies, and the latter because sex has a distinct influence on immune system function^13^. We developed a novel approach to quantify the association of the WGCNA module expression with these two factors (See Online Methods). Figure 9 shows that, in the case of acute HIV infection, most modules are associated with the disease process (**HIVsetB** dataset), but that one module (*purple*) is strongly associated with sex. **Figure S5** shows the module eigengenes for the *yellow* and *purple* modules, demonstrating differential class associations for WGCNA modules; this finding would have been missed had the data not been stratified by both HIV infection status and sex. **Table S4** shows the module statistics for this dataset.

**Figure 9.**
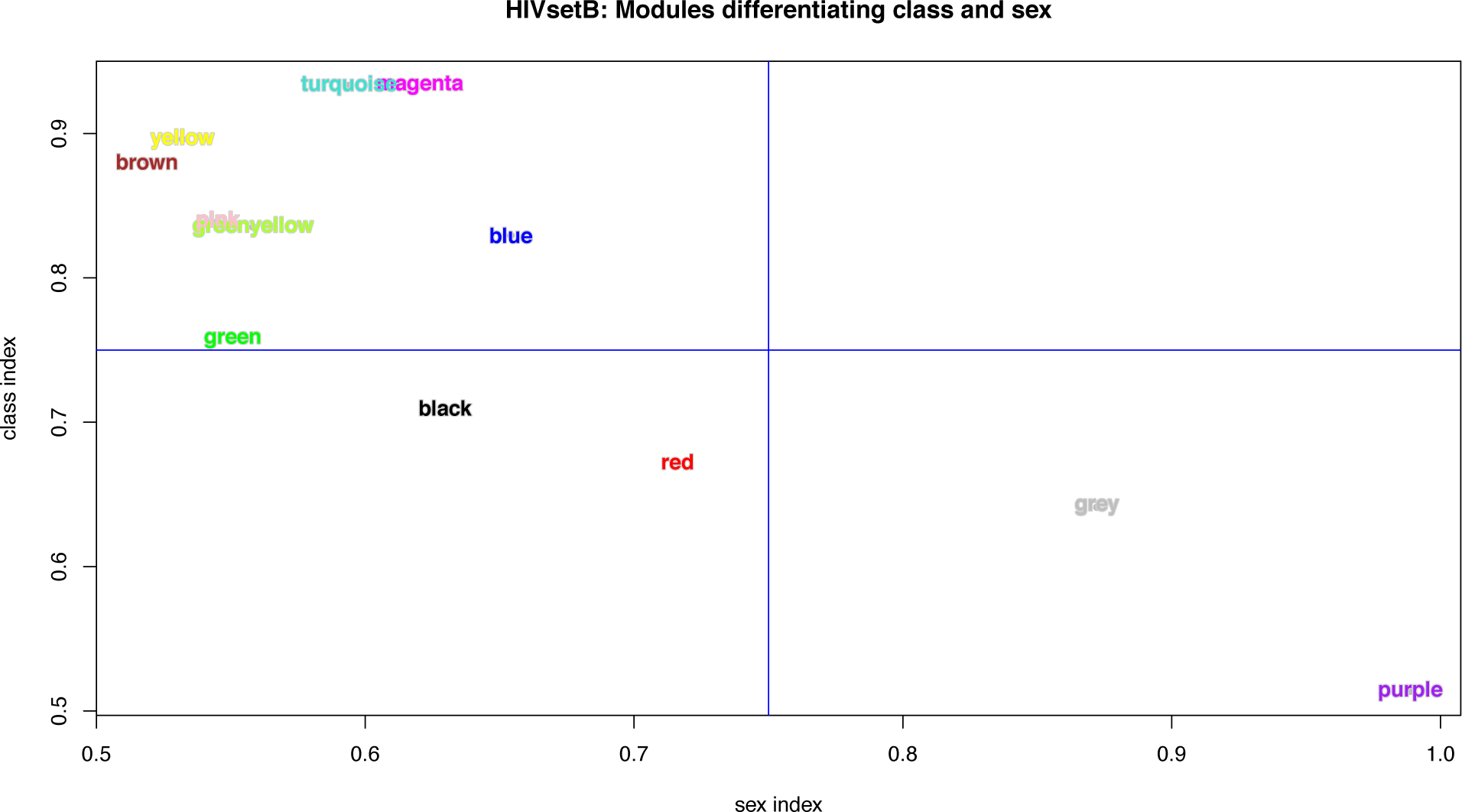
WGCNA module indices. Plot of module indices representing the area-under-the-ROC curve for the two classes for all WGCNA modules in the **HIVsetB** dataset (N_HIV_=30, N_controls_=17, see **Figure S1**). The indices are named per the variable they aim to differentiate (disease class or sex). The class index corresponds to the modAUC1 variable and the sex index corresponds to modAUC2. These indices are calculated form the module eigengenes and given class assignments using functions from the *rocr* package. See text and Online Methods for details.

Additional modules were identified that associated with neither sample class nor sex. These represent biologic processes that manifest in heterogeneity of the sampled population. **Figure S6** plots the study subjects in the HIV infection data set (**HIVsetA**) on two axes represented by two module eigengenes.

### Approach 3: Meta-analysis of multiple datasets

#### Module-level meta-analysis

Meta-analysis of multiple related expression datasets can lead to insights not available from analysis of any single datasets, and can highlight common patterns of transcript abundance across different conditions, or meaningful differences across highly common conditions. We implemented the approach pioneered^11^ and refined^14^ by Chaussabel, *et al* to perform modular transcriptional repertoire analysis^15^ on the six datasets. This approach is particularly suited to meta-analysis, as the composition of the modules is always identical. Figure 10 shows modular patterns for the six datasets in a clustered heatmap as well as the subset of modules with similar expression patterns across the six data sets, demonstrating universal patterns in the immune response to infection. **Supplementary Table S3** lists module functions based on all significant enrichment associations for the modules in Figure 10B. Universal upregulation of interferon-related modules particularly stand out, as does the suppression of modules associated with CD4+ T cells, CD8 T cells and B cells.

**Figure 10.**
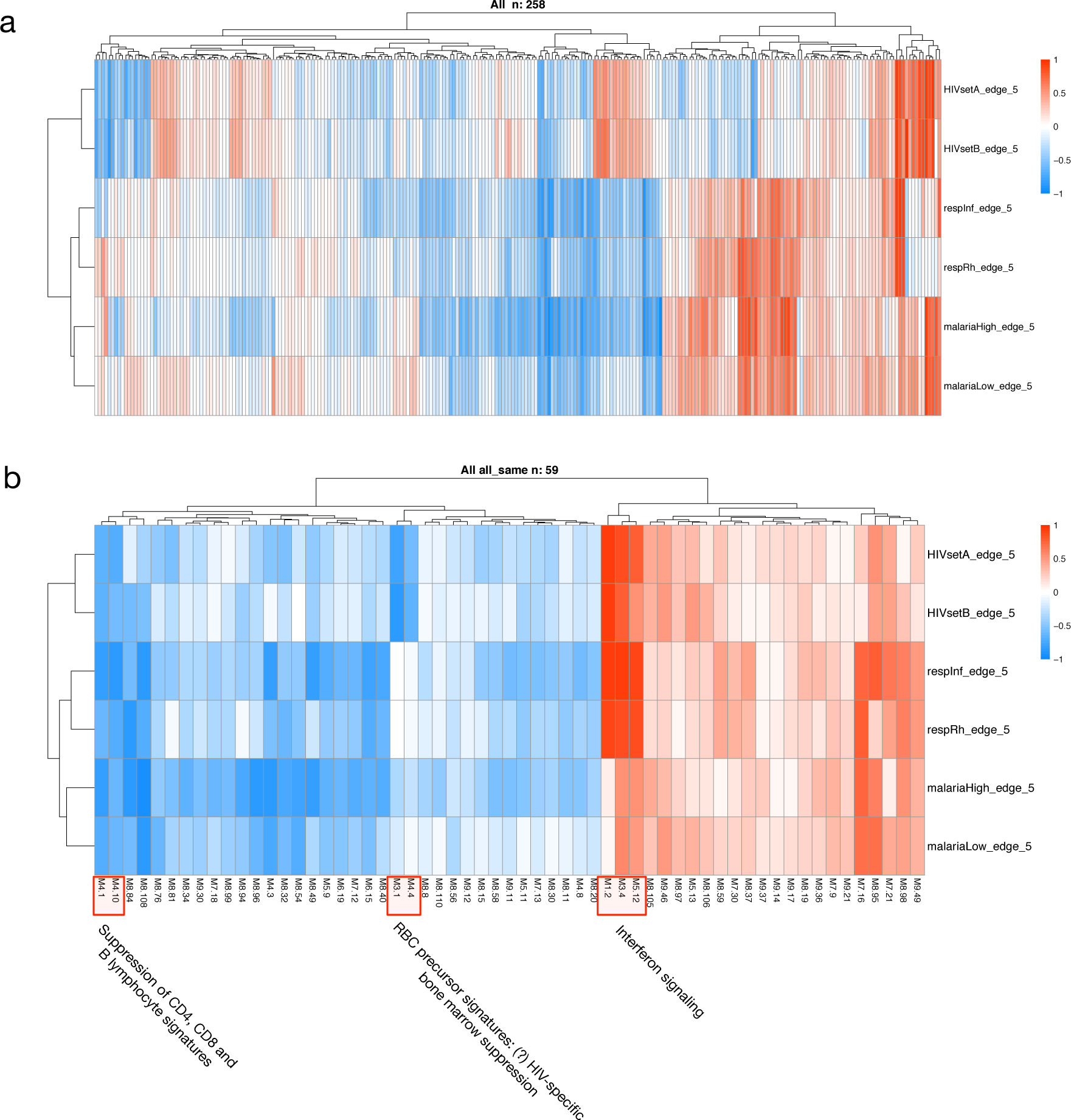
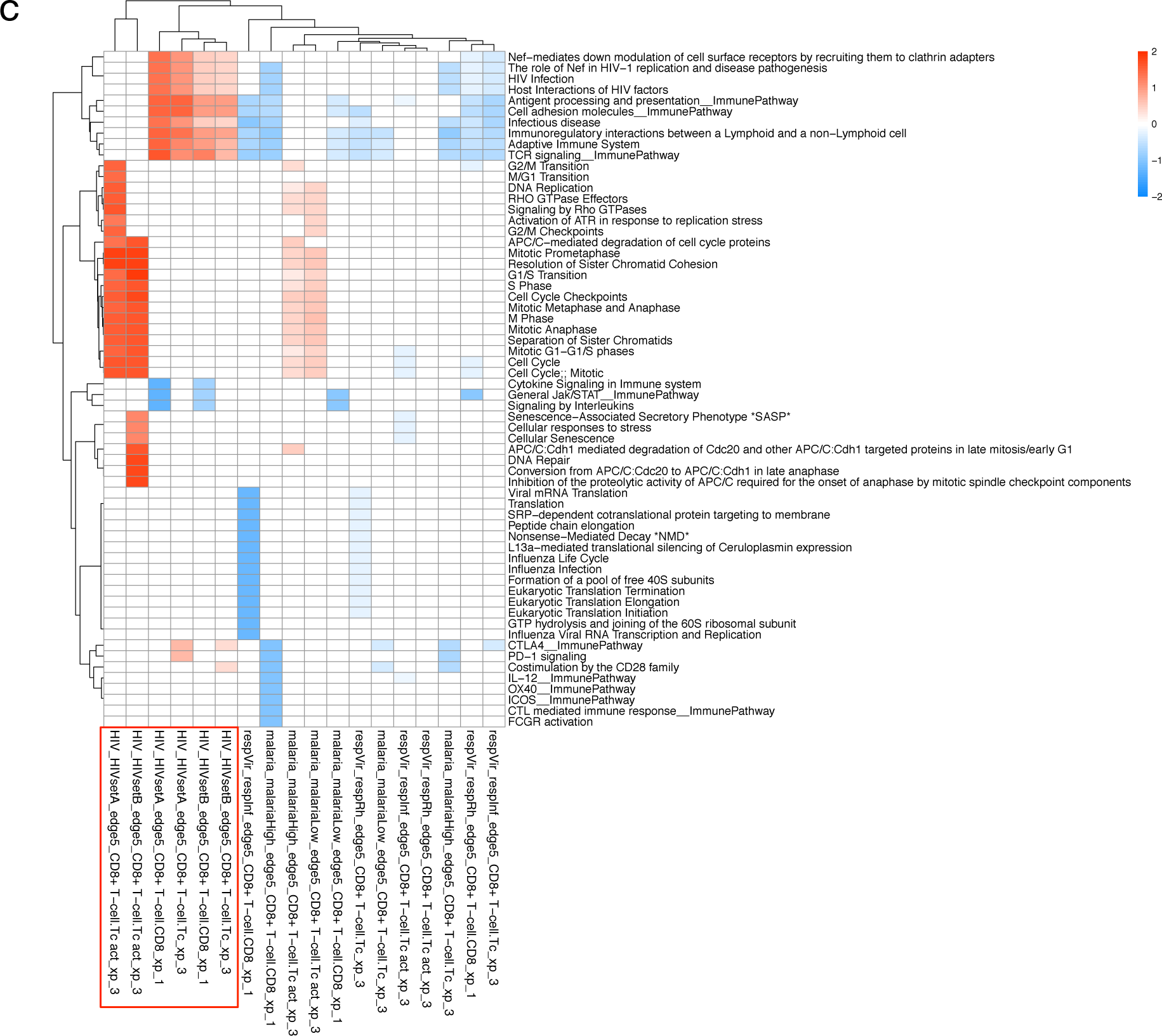
Meta-analysis of transcriptional and cellular patterns. (A) All 258 Chaussabel modules plotted as a heatmap in all six datasets. (B) The subset of modules all expressed in the same direction. Three module groups of interest are identified. (C) Cell/ pathway activity matrix for a single cell type (CD8 + T cell) based on three celltype-gene lists (xp1, xp2, xp3, see Online Methods) in all three conditions. Activity in CD8+ T cells in HIV all cluster together, and differ from both malaria and respiratory infections. Cell labels are constructed by [condition]_[dataset]_[comparison]_[cell class]_[cell type]_[gene list].

#### Meta-analysis at the cell-type level

A second approach to meta-analysis was implemented using virtual cells based on WGCNA modules. Here we compared the pathway scores in a single or several cell types across multiple conditions. For instance, comparing the CD8 T-cell response in acute HIV, acute viral respiratory infection, and symptomatic malaria, we find that proliferation of activated CD8+ T cells characterizes acute HIV infection, and to a lesser extent symptomatic malaria infection. In contrast, there is a suppression of CD8 T cell activity in blood in other conditions, due in part to a reduction in CD8+ T-cell proportions in whole blood (Figure 10C).

## Discussion

Systems immunology aims to understand the complex web of relationships between immune system components (cells, cytokines, effector molecules and other mediators) in immune-mediated disease states. Much progress has been made in single-cell techniques that yield large amounts of information, e.g. single-cell RNAseq and mass cytometry. These approaches are expensive, and not easy to apply in incompletely characterized disease systems, as they require a certain amount of focus (e.g. selection of one or a few cell types for RNAseq, and marker selection for CyTOF). In contrast, RNAseq or microarray analysis of complex tissue samples (like blood) in principle contain information on the transcriptomic state of all cells present in the sample and is thus an unbiased approach. The main difficulty with this lies in the interpretation of the data, and in many cases a complexity-reducing analysis approach is employed, where the focus is placed on differentially expressed genes. Other approaches based on co-expression analysis often fail to explain the drivers of the coexpression patterns.

We demonstrate, using a novel method of aggregating information obtained from clinical and microarray data, an ability to reconstruct many aspects of the immune response, and to discuss this not in the language of probes, genes and signatures, but rather as coordinated biological processes and the cellular context for these processes, allowing the generation of hypotheses at multiple scales. Our multiscale class comparison approach can be used to validate findings from individual papers, for instance the intense NK-cell activation described in influenza^8^.

Using our factorial approach, we can begin to dissect inter individual heterogeneity in transcriptional patterns from transcriptional patterns that are in a causal relationship with defined factors. Application of the two meta-analysis approaches allows comparison of arbitrary datasets to detect similarities and differences at modular and cellular levels. An interesting and somewhat unexpected finding is that acute symptomatic malaria and acute respiratory viral illnesses are more similar to each other than to acute HIV infection, another viral illness. Despite these differences, we demonstrate that, at least for these three rather different infections, a broadly similar pattern of transcriptional module activity can be described.

In summary, ANIMA is both a robust implementation of various well-regarded analytic paradigms in microarray analysis, as well as a framework for integrating these various methods to expose relationships at multiple scales and render these computationally accessible.

## References

1. Ahmed, E. & Hashish, A. H. On modelling the immune system as a complex system. Theory Biosci. 124, 413–418 (2006).

2. Benoist, C., Germain, R. N. & Mathis, D. A plaidoyer for ‘systems immunology’. Immunol. Rev. 210, 229–234 (2006).

3. Mazzocchi, F. Complexity in biology. Exceeding the limits of reductionism and determinism using complexity theory. EMBO Rep. 9, 10–14 (2008).

4. Chaussabel, D., Pascual, V. & Banchereau, J. Assessing the human immune system through blood transcriptomics. BMC Biol. 8, 84 (2010).

5. Langfelder, P. & Horvath, S. WGCNA: an R package for weighted correlation network analysis. BMC Bioinformatics 9, 559 (2008).

6. Chang, H.-H. et al. Transcriptional network predicts viral set point during acute HIV-1 infection. J Am Med Inform Assoc 19, 1103–1109 (2012).

7. Idaghdour, Y. et al. Evidence for additive and interaction effects of host genotype and infection in malaria. Proc Natl Acad Sci USA 109, 16786–16793 (2012).

8. Zhai, Y. et al. Host Transcriptional Response to Influenza and Other Acute Respiratory Viral Infections—A Prospective Cohort Study. PLoS Pathog. 11, e1004869 (2015).

9. Csardi, G. & Nepusz, T. The igraph software package for complex network research. InterJournal (2006).

10. Hardy, G. A. D. et al. Interferon-α is the primary plasma type-I IFN in HIV-1 infection and correlates with immune activation and disease markers. PLoS ONE 8, e56527 (2013).

11. Chaussabel, D. et al. A Modular Analysis Framework for Blood Genomics Studies: Application to Systemic Lupus Erythematosus. Immunity 29, 150–164 (2008).

12. McMichael, A. J., Borrow, P., Tomaras, G. D., Goonetilleke, N. & Haynes, B. F. The immune response during acute HIV-1 infection: clues for vaccine development. Nat. Rev. Immunol. 10, 11–23 (2010).

13. Klein, S. L. & Flanagan, K. L. Sex differences in immune responses. Nat. Rev. Immunol. 16, 626–638 (2016).

14. Obermoser, G. et al. Systems scale interactive exploration reveals quantitative and qualitative differences in response to influenza and pneumococcal vaccines. Immunity 38, 831–844 (2013).

15. Chaussabel, D. & Baldwin, N. Democratizing systems immunology with modular transcriptional repertoire analyses. Nat. Rev. Immunol. 14, 271–280 (2014).

16. Du, P., Kibbe, W. A. & Lin, S. M. lumi: a pipeline for processing Illumina microarray. Bioinformatics 24, 1547–1548 (2008).

17. Barbosa-Morais, N. L. et al. A re-annotation pipeline for Illumina BeadArrays: improving the interpretation of gene expression data. Nucleic Acids Res 38, e17 (2010).

18. Arloth, J., Bader, D. M., Röh, S. & Altmann, A. Re-Annotator: Annotation Pipeline for Microarray Probe Sequences. PLoS ONE 10, e0139516 (2015).

19. Smyth, G. K. Linear Models and Empirical Bayes Methods for Assessing Differential Expression in Microarray Experiments. Statistical Applications in Genetics and Molecular Biology 3, (2004).

20. Gaujoux, R. & Seoighe, C. CellMix: a comprehensive toolbox for gene expression deconvolution. Bioinformatics 29, 2211–2212 (2013).

21. Yu, G. & He, Q.-Y. ReactomePA: an R/Bioconductor package for reactome pathway analysis and visualization. Mol Biosyst 12, 477–479 (2015).

22. Neo4j's Graph Query Language: An Introduction to Cypher. Available at: https://neo4j.com/developer/cypher-query-language/. (Accessed: 3rd January 2017)

